# The BCKDK inhibitor BT2 is a chemical uncoupler that lowers mitochondrial ROS production and *de novo* lipogenesis

**DOI:** 10.1101/2023.08.15.553413

**Authors:** Aracely Acevedo, Anthony E. Jones, Bezawit T. Danna, Rory Turner, Katrina P. Montales, Cristiane Benincá, Karen Reue, Orian S. Shirihai, Linsey Stiles, Martina Wallace, Yibin Wang, Ambre M. Bertholet, Ajit S. Divakaruni

## Abstract

Elevated levels of branched chain amino acids (BCAAs) and branched-chain α-ketoacids (BCKAs) are associated with cardiovascular and metabolic disease, but the molecular mechanisms underlying a putative causal relationship remain unclear. The branched-chain ketoacid dehydrogenase kinase (BCKDK) inhibitor BT2 is often used in preclinical models to increase BCAA oxidation and restore steady-state BCAA and BCKA levels. BT2 administration is protective in various rodent models of heart failure and metabolic disease, but confoundingly, targeted ablation of *Bckdk* in specific tissues does not reproduce the beneficial effects conferred by pharmacologic inhibition. Here we demonstrate that BT2, a lipophilic weak acid, can act as a mitochondrial uncoupler. Measurements of oxygen consumption, mitochondrial membrane potential, and patch-clamp electrophysiology show BT2 increases proton conductance across the mitochondrial inner membrane independently of its inhibitory effect on BCKDK. BT2 is roughly five-fold less potent than the prototypical uncoupler 2,4-dinitrophenol (DNP), and phenocopies DNP in lowering *de novo* lipogenesis and mitochondrial superoxide production. The data suggest the therapeutic efficacy of BT2 may be attributable to the well-documented effects of mitochondrial uncoupling in alleviating cardiovascular and metabolic disease.

## INTRODUCTION

Elevations in circulating levels of the branched-chain amino acids (BCAAs) leucine, isoleucine, and valine are a hallmark of metabolic and cardiovascular disease [1–5]. Early studies dating back over fifty years showed elevated blood levels of BCAAs positively correlated with obesity and blood insulin levels in patients relative to matched lean individuals [6]. Further studies leveraging unbiased metabolomics have reinforced the links between BCAA accumulation and insulin resistance in both humans and pre-clinical rodent disease models [7,8]. In fact, longitudinal studies tracking healthy individuals showed that elevated plasma BCAAs could be predictive of future insulin resistance [8,9].

As with obesity and insulin resistance, epidemiological studies have also linked elevated circulating levels of both BCAAs and their breakdown products, branched-chain α-ketoacids (BCKAs), with cardiovascular disease. In both humans and rodents, several studies point to accumulation of BCAAs and/or BCKAs in heart failure, myocardial infarction (MI) and ischemia, hypertension, and arrhythmia [1,10–14]. Furthermore, elevated plasma BCAAs may be predictive of future obstructive coronary artery disease [15].

As with other metabolites, steady-state levels of BCAAs are a balance of intake and disposal. BCAAs are essential amino acids for humans and animals, meaning they cannot be synthesized *de novo*. Dietary intake is therefore the predominant source of BCAAs, though a minor proportion may arise from the gut microbiome [4]. Along with protein synthesis, BCAA oxidation represents an important sink for clearing BCAAs and maintaining homeostatic levels [4,5].

Oxidation of leucine, isoleucine, and valine is a tightly regulated process. Each BCAA is first deaminated by the branched-chain aminotransferase (BCAT) to its respective BCKA. These BCKAs then undergo irreversible decarboxylation and esterification to their respective R-CoA molecules by the branched-chain α-ketoacid dehydrogenase (BCKDH). They are eventually incorporated into the TCA cycle as acetyl CoA or succinyl CoA. Much like regulation of the pyruvate dehydrogenase by inhibitory kinases and an activating phosphatase, the BCKDH is inhibited by the BCKDH kinase (BCKDK) and activated by the mitochondrial protein phosphatase 2Cm (PP2Cm) [16,17].

The association between increased BCAA and BCKA levels and cardiovascular and metabolic disease is well accepted, but it remains unclear whether these elevated levels have a causative role in disease or represent an indirect epiphenomenon. Several proof-of-concept studies have sought to modify BCAA intake in pre-clinical models to examine how readjusting steady-state levels affects disease pathology. As it relates to insulin resistance, dietary BCAA supplementation has produced mixed results suggesting a complex, contextual effect dependent on the composition of diet and pre-existing health of the animal [4,7]. Dietary restriction of BCAAs, however, reverses insulin resistance in multiple pre-clinical studies [7,18]. Although fewer studies are available examining dietary BCAAs on cardiovascular disease, one report noted oral BCAA administration post-MI impaired contractility and increased infarct size [19].

Numerous studies have manipulated metabolic enzymes to alter steady-state BCAA and BCKA levels and examine the disease-modifying impact. Indeed, whole-body deletion or silencing of the PP2Cm increases circulating BCAA levels and promotes heart failure and worsens ischemia/reperfusion (I/R) injury [12,20]. In line with these results, tissue-specific adenoviral overexpression of the phosphatase lowered BCKA and BCAA levels and prevented ischemic injury in the heart [21] and reduced hepatic triglyceride accumulation and improved hepatic glucose tolerance [22].

Some of the strongest evidence for a causative, pathological relationship between dysregulated BCAA metabolism and cardiometabolic disease comes from pharmacologic inhibition of the BCKDK with BT2. The small molecule BT2 (3,6-dichlorobenzo[b]thiophene-2-carboxylic acid) was designed as a BCKA analog that allosterically binds to the BCKDK [23]. Similar to end-product inhibition of the BCKDK by elevated BCKA levels, the compound relieves kinase inhibition on the BCKDH and promotes BCKA oxidation. In preclinical models of heart failure, systemic BT2 administration lowers cardiac BCAAs and BCKAs and provides cardioprotection in response to both transaortic constriction (TAC) as well as LAD ligation [12,19–21]. In various rodent models of cardiometabolic disease, BT2 lowers circulating BCAA and BCKAs, improves glucose tolerance and insulin sensitivity, and reduces *de novo* lipogenesis [22,24–26].

Remarkably, however, the recent development of tissue-specific mouse models has complicated our understanding of how dysregulated BCAA metabolism may relate to cardiovascular and metabolic disease. Cardiac-specific loss of BCKDK does not reproduce the protective effects in either hemodynamic or ischemic models of heart failure [27]. It could be argued that the proportion of BCAA oxidation in the heart is too low for the cardiac-specific knockout to have an effect on circulating BCAA levels similar to BT2 [28]. However, inducible skeletal muscle-specific *Bckdk* ablation also failed to elicit cardioprotection despite lowering cardiac BCAAs and plasma BCAAs and BCKAs [27]. Furthermore, tissue specific-loss of BCKDK in skeletal muscle does not reproduce the effects of BT2 in improving insulin sensitivity despite lowering plasma BCAAs and BCKAs [29]. Similar loss-of-function studies show that hepatic BCKDK loss also fails to improve insulin sensitivity [26,29], though the liver has less apparent control over circulating BCAA levels than muscle. While the results could suggest that whole-body BCKDK inhibition cannot be reproduced by *Bckdk* ablation in individual tissues, they perhaps also suggest an alternative mechanism by which BT2 confers protection from heart failure and metabolic disease distinct from BCAA oxidation.

Here we present evidence that BT2 is a mitochondrial uncoupler. Measurements of respiration, mitochondrial membrane potential, and inner membrane electrophysiology all demonstrate that BT2 – a lipophilic weak acid – is itself a chemical uncoupler. BT2 also phenocopies the prototypical uncoupler 2,4-dinitrophenol (DNP) in lowering mitochondrial H_2_O_2_ efflux and reducing *de novo* lipogenesis. Importantly, chemical uncoupling of mitochondria provides a single, unified mechanism by which BT2 could plausibly confer protection against heart failure (lowering mitochondrial ROS production) and cardiometabolic disease (increasing energy expenditure) independent of effects on steady-state BCAA levels.

## MATERIALS AND METHODS

### Animals

All animal protocols and procedures were approved and performed in accordance with the NIH Guide for the Care and Use of Laboratory Animals and the UCLA Animal Research Committee (ARC). C57BL/6J male mice aged 8-12 weeks were purchased from The Jackson Laboratory. Male Sprague Dawley rats aged between 7-10 weeks (∼200-300 g) were purchased from Envigo.

### Reagents

BT2 (3,6-dichlorobenzo[b]thiophene-2-carboxylic acid) was purchased from MedChemExpress (#HY114855) and all other uncouplers were purchased from Sigma-Aldrich [FCCP (Carbonyl cyanide 4-trifluoromethoxyphenylhydrazine; #C2920), DNP (2,4-dinitrophenol; #42195), Bam15 (#SML-1760)]. Stocks were made and stored at -20°C at the following concentrations: BT2 (40 mM and 400 mM in DMSO); FCCP (10 mM in 95% ethanol); DNP (20 mM in DMSO); Bam15 (40 mM in DMSO).

### Cell culture

#### Cardiomyocytes

Neonatal rat ventricular myocytes (NRVMs) were isolated from post-natal P1-P3-day old Sprague Dawley rat pups of mixed sex as previously described [30,31]. Cells were plated onto Agilent Seahorse XF96 cell culture plates or 12-well cell culture dishes coated with 0.1% gelatin (Sigma #G1393) in DMEM/F12 medium (Gibco #11330057) supplemented with 10% (v/v) FBS, 100 U/ml penicillin, and 100 μg/mL streptomycin. After 24 hr., medium was changed to DMEM/F12 lacking FBS but with antibiotics as before. Human induced pluripotent stem cell (iPSC) cardiomyocytes (iCell cardiomyocytes; Fujifilm Cellular Dynamics International #01434) were maintained according to manufacturer’s guidelines. Cells were grown and maintained in a humidified 5% CO_2_ incubator at 37°C.

#### 3T3-L1 adipocytes

3T3-L1 pre-adipocytes were maintained below 70% confluency in DMEM (Gibco #11965) supplemented with 10% (v/v) bovine calf serum (ATCC 30-2020), 100U/mL penicillin, 100 μg/mL streptomycin, and 1 mM sodium pyruvate. Cells were plated onto XF96 cell culture plates or 12-well tissue culture dishes and allowed to grow for 48 hr. (plating on Day -2). On Day 0, differentiation was begun by changing the medium to maintenance medium [DMEM supplemented with 10% (v/v) FBS, 100U/mL penicillin, 100 μg/mL streptomycin, 1 mM sodium pyruvate, 10 mM HEPES] supplemented with 1 μg/mL insulin (Sigma #I0515), 0.25 μM dexamethasone (Sigma #D4904), 0.5 mM methylisobutylxanthine (IBMX, Sigma I5879), and 100 nM rosiglitazone (Sigma #R2408). On Day 2, medium was changed to maintenance medium supplemented only with 1 μg/mL insulin, and on Days 4 and 6 medium was replaced to maintenance medium with no further additions. Cells were treated with compound on Day 8 as described elsewhere in the Materials and Methods.

#### C2C12 myoblasts

C2C12 myoblasts were maintained in DMEM (Gibco #11965) supplemented with 10% (v/v) FBS, 100U/mL penicillin, 100 μg/mL streptomycin, and 1 mM sodium pyruvate. Cells with genetic ablation of *Slc25a4* and *Slc25a5* (encoding AAC1 and AAC2) were generated and described previously [32].

### Mitochondrial Isolation

Mitochondrial isolation from rodent heart and liver was conducted according to well established protocols [33] and all steps were conducted on ice or at 4°C. Mitochondrial protein content was measured by the bicinchoninic acid (BCA) assay.

#### Liver mitochondria

Mouse or rat livers were cleaned, minced, and drained of blood in ice-cold MSHE [210 mannitol, 70 mM sucrose, 5 mM HEPES, 1 mM EGTA, and 0.2% (w/v) fatty acid-free BSA] at pH 7.2 at 4°C. Livers were disrupted with using 2 strokes of a drill-driven Teflon-on-glass Dounce homogenizer with roughly 1mL of buffer for every 100 mg of tissue. The homogenate was spun at 12,000*g* for 10 minutes at 4°C to remove any contaminating fat. The pellet was resuspended and centrifuged at 800*g* for 5 minutes at 4°C to remove debris. The supernatant was filtered through two layers of wet cheesecloth and centrifuged at 12,000*g* for 10 minutes at 4°C. The light, ‘fluffy’ layer of the pellet was removed, and the mitochondrial pellet was resuspended and centrifuged again at 12,000*g* at 4°C. For the third and final centrifugation step, the pellet was washed and resuspended in MSHE lacking BSA and centrifuged again at 4°C. The final mitochondrial pellet was resuspended in MSHE lacking BSA and kept at a concentration greater than 100 mg/mL and kept on ice.

#### Heart mitochondria

Mouse or rat hearts were quickly removed from the euthanized animal while still beating and minced, cleaned of blood, and homogenized using a hand-held tissue disruptor (IKA Ultra-Turrax) in ice-cold MSHE. The homogenate was centrifuged at 900*g* for 10 minutes at 4°C. The supernatant was then centrifuged at 9,000*g* for 10 minutes at 4°C, and the remaining pellet was washed and re-centrifuged at 9,000*g* at 4°C in medium lacking BSA. The final mitochondrial pellet was resuspended in MSHE lacking BSA and kept at a concentration greater than 25 mg/mL and kept on ice.

### Respirometry

All oxygen consumption measurements were conducted using an Agilent Seahorse XF96 or XF^e^96 Analyzer. Experiments were conducted at 37°C and at pH 7.4 (intact cells) or 7.2 (isolated mitochondria and permeabilized cells). For experiments with intact or permeabilized cells, only the inner 60 wells were used and the outer rim was filled with 200 μL of PBS throughout the incubation to minimize variance in temperature and evaporative effects across the plate. All respiratory parameters were corrected for non-mitochondrial respiration and background signal from the instrument with addition of 200 nM rotenone and 1 μM antimycin A and calculated according to well established best practices [34,35].

#### Intact cells

Respiration was measured in NRVMs (4.0x10^4^ cells/well), iPSC cardiomyocytes (1.5x10^4^ cells/well), C2C12 myoblasts (1.5x10^4^ cells/well), and differentiated 3T3-L1 adipocytes (2.5x10^3^ cells/well) in DMEM assay medium composed of DMEM (Sigma #5030) supplemented with 31.6 mM NaCl, 3 mg/L phenol red, 5 mM HEPES, 10 mM glucose, 2 mM glutamine, and 2 mM pyruvate. Where appropriate, cells were offered oligomycin (2 μM) and maximal respiration was estimated with 750 nM FCCP.

#### Permeabilized cells

Cells were permeabilized with 3 nM recombinant perfringolysin O (PFO) and assayed as previously described [36] in MAS buffer [220 mM mannitol, 70 mM sucrose, 10 mM KH_2_PO_4_ 5 mM MgCl_2_, 2 mM HEPES (pH 7.2 at 37°C), 1 mM EGTA, 0.2% (w/v) BSA]. Where indicated medium was supplemented with 4 mM ADP and either 10 mM pyruvate with 1 mM malate or 10 mM succinate with 2 μM rotenone. Where indicated, oligomycin was used at 2 μM.

#### Isolated mitochondria

Oxygen consumption rates in isolated mitochondria were measured in MAS buffer described previously according to well established protocols [36,37]. As indicated in the figure legends, BSA was either omitted from the assay medium or used at 0.001% (w/v) to avoid sequestering lipophilic compounds under investigation. Where indicated, MAS buffer was supplemented with 10 mM pyruvate with 1 mM malate (Pyr/Mal), 5 mM glutamate with 5 mM malate (Glu/Mal), or 10 mM succinate with 2 μM rotenone (Succ/Rot). Mitochondria were offered oligomycin at 2 μM and FCCP at 1 μM where indicated.

#### Normalization

All intact and permeabilized cell oxygen consumption rates were normalized to cell number. Cells were immediately fixed after the assay with 2% (v/v) paraformaldehyde in PBS and stored at 4°C for up to 14 days. Nuclei were stained with Hoescht (ThermoFisher #33342) overnight at 4°C and quantified using the Operetta High Content Imaging System (Perkin Elmer). All oxygen consumption rates in isolated mitochondria were normalized to microgram of total mitochondrial protein in the microplate well unless otherwise indicated.

### Mitochondrial Membrane Potential

#### Isolated mitochondria

The mitochondrial membrane potential was measured using the quenched fluorescence using a high concentration of tetramethylrhodamine, ethyl ester (TMRE) as has been previously described [38]. TMRE fluorescence will self-quench upon accumulation in the mitochondrial matrix at high concentrations. As such, lowering the membrane potential will decrease dye uptake and self-quenching, thereby increasing the observed fluorescent signal. Mitochondria isolated from rat liver or rat heart (0.6 mg/mL) were incubated in MAS buffer supplemented with 0.001% BSA, 4 mM ADP, 5 mM succinate, 2 µM rotenone, and 5 µM TMRE at 37°C for 10 minutes in a black-walled 96-well microplate shielded from light. After incubation, fluorescence was measured (549ex./575em.) using a Tecan Spark multimode plate reader. Compounds under investigation were added immediately prior to the 10 min. incubation period.

#### Cardiomyocytes

The mitochondrial membrane potential in cells was measured with TMRE using the Image Xpress Micro Confocal high-content imaging system (Molecular Devices). NRVMs and iPSC-derived iCell cardiomyocytes were plated onto collagen-coated, black-walled PhenoPlates (Perkin-Elmer) at either 6.0x10^4^ (NRVMs) or 1.5x10^4^ (iCell) cells/well and maintained as described earlier in the Materials and Methods. 75 minutes prior to conducting measurements, medium was exchanged for DMEM lacking glucose, phenol red, and sodium bicarbonate (Sigma #5030) supplemented with 31.6 mM NaCl, 10 mM glucose, 2 mM glutamine, 2 mM pyruvate, and 5 mM HEPES along with 10 nM TMRE (Invitrogen #T669) and 200 nM MitoTracker Green FM (Catalog # M7514). After allowing the dyes to equilibrate for 1 hr., BT2 (80 µM), DNP (10 µM) and FCCP (1 µM) were added, and measurements of the first microplate well began after 15 minutes. Circularity is defined with “1” being a perfect circle and “0” a straight line.

### Patch-clamp recordings

Proton conductance across the mitochondrial inner membrane was conducted on mitoplasts derived from mouse heart mitochondria as previously described [32,39]. Patch-clamp recording was performed from isolated heart mitoplasts of mice. The mitoplasts used for patch-clamp experiments were 3–5 μm in diameter and typically had membrane capacitances of 0.3–1.2 pF. Both the bath and pipette solutions were formulated to record H^+^ currents and contained only salts that dissociate into large anions and cations that are normally impermeant through ion channels or transporters. A low pH gradient across the IMM was used (pH 7.5 and 7.0 on the matrix and cytosolic sides, respectively).

Pipettes were filled with 130 mM tetramethylammonium hydroxide (TMA), 1 mM EGTA, 2 mM Tris-HCl, and 100 mM HEPES. pH was adjusted to 7.5 with D-gluconic acid, and tonicity was adjusted to ∼360 mmol/kg with sucrose. Typically, pipettes had resistances of 25–35 MΩ, and the access resistance was 40–75 MΩ. Whole-mitoplast *I*_H_ was recorded in the bath solution containing 100 mM HEPES and 1 mM EGTA (pH adjusted to 7 with Trizma base, and tonicity adjusted to ∼300 mmol/kg with sucrose). All experiments were performed under continuous perfusion of the bath solution. All electrophysiological data presented were acquired at 10 kHz and filtered at 1 kHz.

### Hydrogen peroxide efflux

H_2_O_2_ efflux was measured in isolated mitochondria as described previously [40] using a Tecan Spark multimode plate reader and an Amplex Red-based detection system. Mitochondria (2 mg/mL) were incubated in SHE buffer (250 mM sucrose, 10 mM HEPES, 1 mM EGTA, pH 7.2 at 37°C) supplemented with 2.5 μM Amplex Red (ThermoFisher #A12222) and 5 U/mL horseradish peroxidase (ThermoFisher #31491). 10 mM succinate was added to the incubation, and the rate of H_2_O_2_ was measured over 2-3 minutes. The signal was calibrated to known amounts of H_2_O_2_ added on top of isolated mitochondria. Compounds under investigation were pre-incubated with mitochondria in SHE buffer for 2-5 minutes prior to the addition of succinate.

### Stable isotope tracing and *de novo* lipogenesis

#### Cell culture

3T3-L1 pre-adipocytes were seeded at 2.5x10^4^ cells/well in 12-well dishes and differentiated as described earlier in the Materials and Methods. On Day 8, cells were changed into maintenance medium (made with DMEM A1443001) with the following changes: 2% (v/v) FBS rather than 10% to reduce protein binding of compounds under investigation, and uniformly labeled [^13^C_6_]-glucose (Cambridge Isotope Laboratories #CLM-1396) was used instead of ^12^C unlabeled glucose. BT2 (120 µM) and DNP (40 µM) were added during this step and used at higher concentrations than in other experiments to account for binding to albumin in 2% (v/v) FBS. After 72 hr. cells were extracted for GC/MS analysis as described below. A matched 12-well dish was used for normalization of metabolite levels to cell number. On the day of the extraction, cells were fixed with 2% (v/v) paraformaldehyde in PBS and stored at 4°C for no less than 7 days. Nuclei were stained with Hoescht (ThermoFisher #33342) overnight at 4°C and quantified using the Operetta High Content Imaging System (Perkin Elmer).

Neonatal rat ventricular myocytes were seeded at 4.0x10^5^ cells/well in 12-well dishes coated with 0.1% (w/v) gelatin in DMEM/F12 medium described earlier. After 24 hr. medium was exchanged into custom DMEM formulated without glucose, glutamine, or leucine (Sciencell Laboratories) and supplemented with 10 mM glucose, 2 mM GlutaMAX, or 0.8 mM leucine, with a uniformly labeled [^13^C_6_] label on either glucose or leucine (#CLM-2262). After 24 hr. cells were extracted for GC/MS analysis as described below.

#### Derivatization and mass spectrometry

Cell preparation and stable isotope tracing measuring incorporation of isotopic labels into polar metabolites and palmitate was conducted as previously described [41,42]. Metabolite extraction was conducted with a Folch-like extraction with a 5:2:5 ratio of methanol:water:chloroform. 12-well dishes were kept on ice and quickly washed with ice-cold 0.9% (w/v) NaCl. Cells were then scraped in ice-cold methanol and water containing 5 µg/mL norvaline (Sigma #N7502), an internal standard. Chloroform containing 20 µM [U-^2^H_31_]-palmitate (Cambridge Isotope Laboratories #DLM-215) as an internal standard was then added to the samples. Samples were then vortexed for 1 min and centrifuged at 10,000g for 5 min at 4°C.

The polar fraction (top layer) was removed and the samples were dried overnight using a refrigerated CentriVap vacuum concentrator (LabConco). Metabolites (50 nmol to 23 pmol) were extracted alongside the cell samples to ensure the signal fell within the linear detection range of the instrument. The dried polar metabolites were reconstituted in 20 µL of 2% (w/v) methoxyamine in pyridine prior to a 45-min incubation at 37°C. Subsequently, 20 µL of MTBSTFA with 1% tert-butyldimethylchlorosilane was added to samples, followed by an additional 45-min incubation at 37°C. Samples were analyzed using Agilent MassHunter software and FluxFix software (http://fluxfix.science) was used to correct for the abundance of natural heavy isotopes against an in-house reference set of unlabeled metabolite standards [43].

The lower, organic fraction was dried under air and then solubilized in 500 µL of acidified methanol [2% (v/v) H_2_SO_4_ in methanol] for 2 hr. at 50°C to generate fatty acid methyl esters (FAMEs). After this incubation, 100 µL of saturated NaCl was added, followed by 500 µL of hexane. This mixture was vortexed for 2 min and the upper hexane layer was collected in a new microfuge tube. An additional 500 µL of hexane was added to the original MeOH/NaCl tube and the process repeated to collect any residual FAMEs not obtained by the first addition. The pooled hexane extracts were then dried under airflow and resuspended in 75 µL of hexane. FAMEs were analyzed by GC-MS analysis. *De novo* lipid synthesis was estimated by isotopomer spectral analysis (ISA) [42,44] and calculated using MATLAB software.

Samples were analyzed using a DB-35 column (Agilent Technologies). Information regarding additional technical specifications is available elsewhere [41,42].

### qPCR

3T3-L1 adipocytes were plated, maintained, and treated with either BT2 or DNP in 12-well dishes identically to those used for GC/MS analysis. Transcript levels were measured using quantitative polymerase chain reaction (qPCR). RNA was extracted using the RNeasy Mini Kit (Qiagen, 74106), and cDNA was generated using the High-Capacity cDNA Reverse Transcription Kit (Applied Biosystems, 4368814). The PowerUp SYBR Green Master Mix kit (Applied Biosystems, A25743) and a QuantStudio 5 (Applied Biosystems) were used for qPCR analysis. Relative gene expression was calculated using the 1ΔΔC_t_ method with *36b4* as a reference gene.

### Statistical analysis

All statistical parameters, including the number of replicates (n), can be found in the figure legends. Statistical analyses were performed using Graph Pad Prism 5 software. Data are presented as the mean ± SEM unless otherwise specified. Individual pairwise comparisons were performed using two-tailed Student’s t-test. For experiments involving two or more groups, data were analyzed by one-way, repeated measures ANOVA followed by Dunnett’s post-hoc multiple comparisons tests against vehicle controls. Data were assumed to follow a normal distribution (no tests were performed). Values denoted as follows were considered significant: *, p < 0.05; **, p < 0.01; ***, p < 0.001.

## RESULTS

### BT2 uncouples mitochondria in rodent and human cardiomyocytes

To identify uncharacterized metabolic targets or mechanisms of BT2 action independent of its inhibitory effect on the BCKDK, we measured the effect of BT2 on the oxygen consumption rate in neonatal rat ventricular myocytes (NRVMs). Acute BT2 administration 5 minutes prior to initial measurements revealed a consistent increase in the basal respiration rate (**Figs. 1A, B**). This increase persisted in the presence of the ATP synthase inhibitor oligomycin, but no change was observed when measuring the maximal respiratory capacity with FCCP. The result mimics the hallmark profile of mitochondrial uncoupling [34,45]: BT2 increased mitochondrial oxygen consumption independently of (or ‘uncoupled’ from) ATP synthesis. Indeed, the uncoupling response was preserved when cardiomyocytes were stimulated with norepinephrine (**Fig. 1B**). Moreover, acute BT2 administration also increased respiration associated with proton leak in iCell human iPSC-derived cardiomyocytes both with and without adrenergic activation (**Fig. 1C**). Given that BT2 acutely increases oligomycin-insensitive respiration in human and rodent cardiomyocytes, we hypothesized that it can act as a chemical uncoupler.

**Figure 1.**
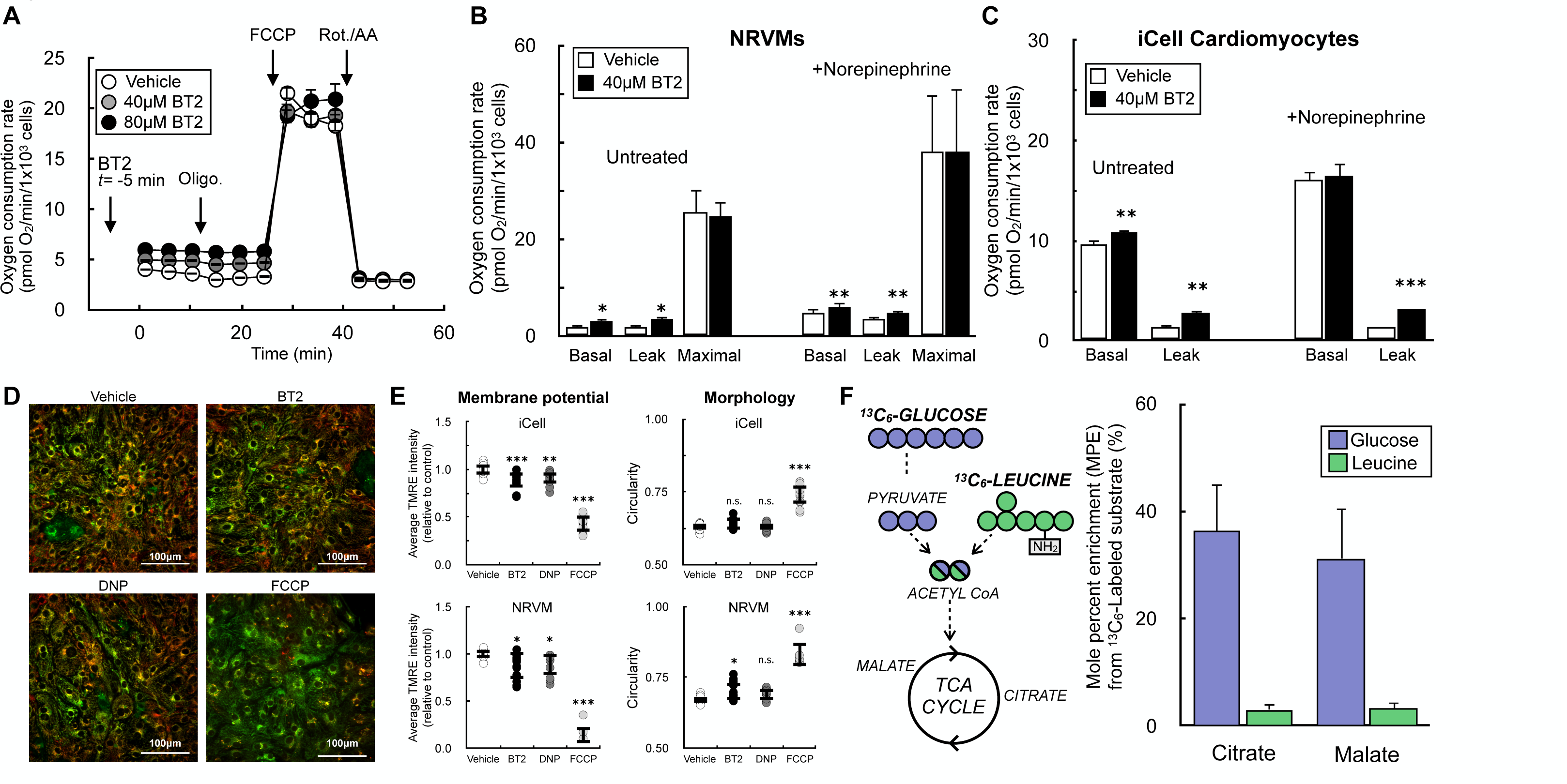
BT2 acutely uncouples mitochondria in rat and human cardiomyocytes. (A) Representative oxygen consumption trace of neonatal rat ventricular myocytes (NRVMs) offered 40μM BT2, 80μM BT2, or vehicle control (DMSO) 5 min prior to conducting measurements. 40μM and 80μM BT2 were chosen as concentrations because they are 10-fold above the EC_50_ (40μM) or previously used in the literature (80μM) [83]. (n=10 technical replicates from a single experiment). (B) Collated oxygen consumption rate parameters for NRVMs offered 40μM BT2 in the presence or absence of 1μM norepinephrine. (n=4 biological replicates). (C) Collated oxygen consumption rate parameters for human iPSC-derived iCell cardiomyocytes offered 40μM BT2 in the presence or absence of 1μM norepinephrine as in (B). (n=4 biological replicates). (D) Representative images for iCell cardiomyocytes treated for 30-45 min. with 80μM BT2, 10μM DNP, 1μM FCCP, or vehicle control. (E) (*Left)* Average TMRE intensity relative to vehicle control for treatments as in (D) with iCell cardiomyocytes (top) and NRVMs (bottom). (*Right*) Circularity as a measure of mitochondrial morphology for treatments as in (D) with iCell cardiomyocytes (top) and NRVMs (bottom). (n=7-15 technical replicates collated from n=3 biological replicates for each cell type). (F) (*Left*) Simplified schematic of uniformly labeled ^13^C_6_-glucose or ^13^C_6_-leucine enriching TCA cycle intermediates. (*Right*) Mole percent enrichment (M.P.E.) of TCA cycle intermediates citrate and malate in NRVMs with the ^13^C label provided on either glucose or leucine. (n=3 biological replicates). All data are mean ± S.E.M. Statistical analysis was conducted with a pairwise Student’s t test [(B) and (C)] or ANOVA followed by Dunnett’s post-hoc multiple comparisons tests [(E)] as appropriate. *, p < 0.05; **, p < 0.01; ***, p < 0.001.

Chemical uncouplers such as FCCP or 2,4-dinitrophenol (DNP) increase the oxygen consumption rate because they stimulate consumption of the mitochondrial membrane potential independently of ATP synthesis, thereby causing increased activity of the respiratory chain [45]. We therefore measured the resting mitochondrial membrane potential in iCell cardiomyocytes and NRVMs in response to acute, 30-45 min. treatment with BT2 (**Figs. 1D, E**). In both model systems, fluorescence of the lipophilic cation dye TMRE showed a ∼10% decrease in the steady-state membrane potential with little change to mitochondrial morphology in response to BT2. This result could be reproduced by the classic uncoupler DNP. Moreover, the changes observed with BT2 and DNP were qualitatively, but not quantitatively, matched by the extreme changes in membrane potential observed from the highly potent uncoupler FCCP. The demonstration that BT2 slightly lowers the steady-state membrane potential similarly to DNP aligns with the oxygen consumption results and supports the hypothesis that BT2 can act as a chemical uncoupler.

We further sought to determine whether our *in vitro* cardiomyocyte models behaved similarly to recent *in vivo* human and rodent studies that show or infer little BCAA oxidation in the heart relative to other substrates [27,28]. We offered NRVMs medium with a uniform ^13^C_6_ label on either glucose or leucine to measure relative incorporation of each substrate into the TCA cycle (**Fig. 1F**). Incorporation of labeled leucine into the TCA cycle intermediates citrate and malate were more than 10-fold lower than incorporation of labeled glucose. The lack of robust rates of BCAA oxidation in this system – in addition to the acute time-frame in which BT2 increases in proton leak-associated respiration – suggested a mechanism independent of BCKDK inhibition. As such, we sought to confirm this by using reductionist systems where specific metabolic pathways and reactions could be isolated.

### BT2 is a chemical uncoupler

To formally rule out the involvement of the BCKDK in the uncoupling phenotype observed in rat and human cardiomyocytes, we measured the acute effects of the compound on isolated mitochondria. Isolated mitochondria are a well-defined system that can be used to isolate specific metabolic reactions and pathways [34,46]. We first measured respiration in mitochondria offered specific substrates to isolate the effects of BT2 on targeted metabolic pathways that bypass the BCKDK. In isolated liver mitochondria offered pyruvate and malate, acute BT2 treatment 5 minutes prior to measurements elicited a similar, specific effect on mitochondrial uncoupling as we had observed in intact cells. BT2 caused a concentration-dependent increase in respiration associated with proton leak (‘State 4_o_’), but no change was observed in ADP-stimulated (‘State 3’) or FCCP-stimulated (‘State 3_u_’) respiration (**Figs. 2A, B**). In liver mitochondria isolated from both rat and mouse, BT2 showed a similar concentration-dependent increase in State 4_o_ respiration irrespective of whether the mitochondria were provided pyruvate, glutamate, or succinate as a respiratory substrate (**Fig. 2C**). Depending on the substrate and species, an increase in proton leak-associated respiration was observed at BT2 concentrations as low as 2.5 μM. In mitochondria isolated from rat and mouse hearts, BT2 also acutely increased State 4_o_ respiration independent of substrate provided to the mitochondria (**Fig. 2D**). In total, respiration with isolated mitochondria showed that BT2 can uncouple isolated mitochondria at single-digit micromolar concentrations regardless of the respiratory substrate provided.

**Figure 2.**
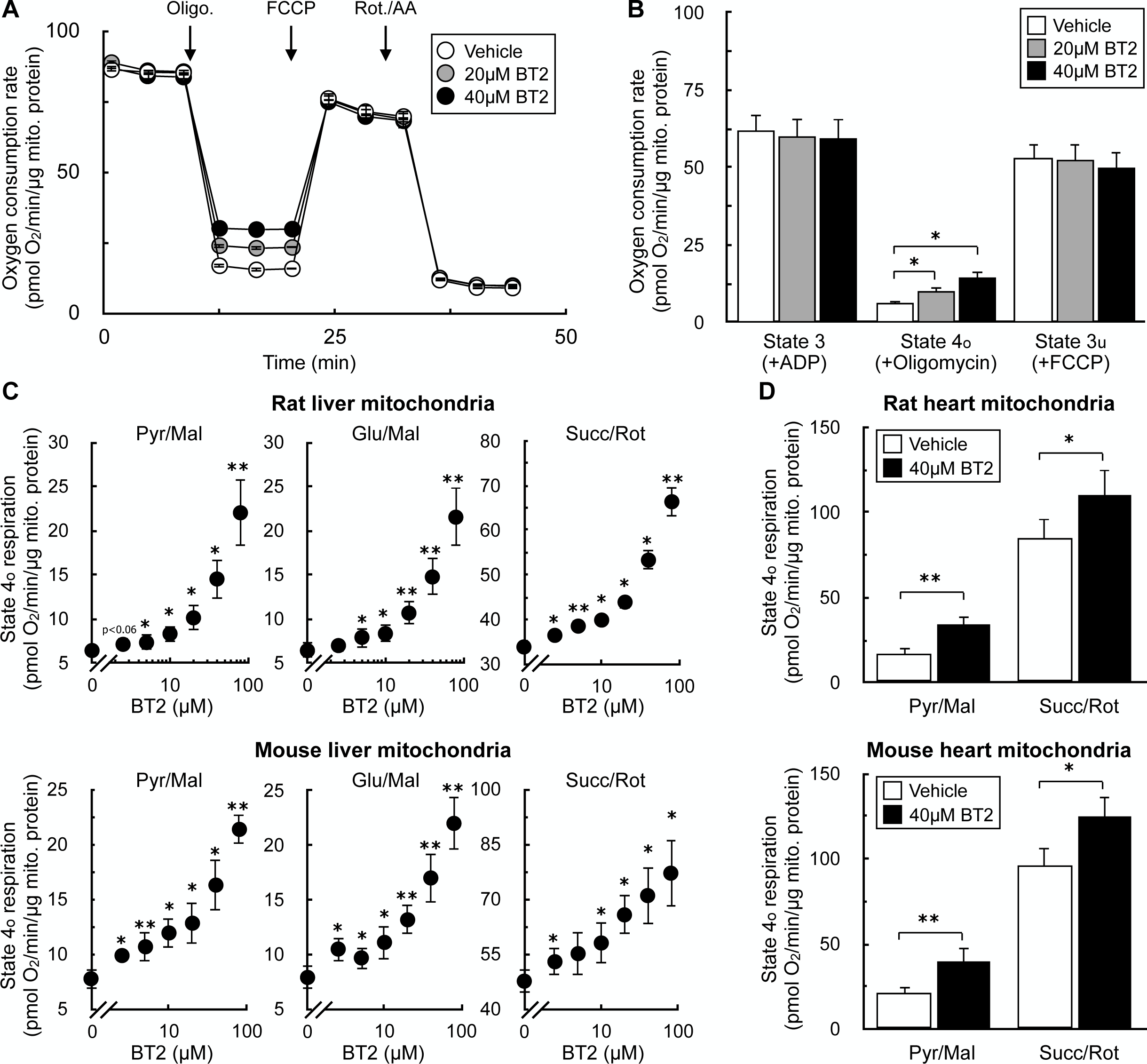
BT2 increases State 4_o_ respiration in isolated mitochondria. (A) Representative oxygen consumption trace of isolated liver mitochondria acutely treated with vehicle control, 20μM BT2, or 40μM BT2 5 min prior to initial measurements. Mitochondria were offered pyruvate, malate, and ADP in the experimental medium and respiration was measured in response to sequential injections of oligomycin, FCCP, and rotenone with antimycin A. (n=5 technical replicates from a single experiment). (B) Collated oxygen consumption rate parameters for isolated liver mitochondria as in (A). (n=4 biological replicates). (C) State 4_o_ respiration in isolated mitochondria from rat liver (*top*) or mouse liver (*bottom*). Experiments conducted as in (A) with pyruvate/malate (*left*), glutamate/malate (*middle*), or succinate/rotenone (*right*) offered as respiratory substrates. [BT2] = 2.5, 5, 10, 20, 40, and 80μM. (n=4 biological replicates). (D) State 4_o_ respiration in isolated mitochondria from rat heart (*top*) or mouse heart (*bottom*) acutely offered either vehicle control (DMSO) or 40μM BT2 5 min prior to initial measurements. Experiments conducted as in (A) with either pyruvate/malate (*left*) or succinate/rotenone (*right*) offered as respiratory substrates. All data are mean ± S.E.M. Statistical analysis was conducted with a pairwise Student’s t test [(D)] or ANOVA followed by Dunnett’s post-hoc multiple comparisons tests [(B) and (C)] as appropriate. *, p < 0.05; **, p < 0.01.

As in Figure 1, we sought to corroborate our oxygen consumption results with measurements of the mitochondrial membrane potential. We therefore examined whether acute BT2 administration would lower the membrane potential in isolated mitochondria by measuring the loss of quenched TMRE fluorescence [47]. The approach was validated by examining the acute effects of oligomycin and FCCP: ATP synthase inhibition by oligomycin increased the membrane potential and further quenched the TMRE signal, whereas the potent uncoupler FCCP caused a substantial loss of the quenched fluorescence (**Fig. 3A**). As expected, acute BT2 treatment lowered the quenched fluorescence of TMRE in both heart and liver mitochondria, indicating a reduced membrane potential and further suggesting the compound itself is a chemical uncoupler (**Fig. 3B**).

**Figure 3.**
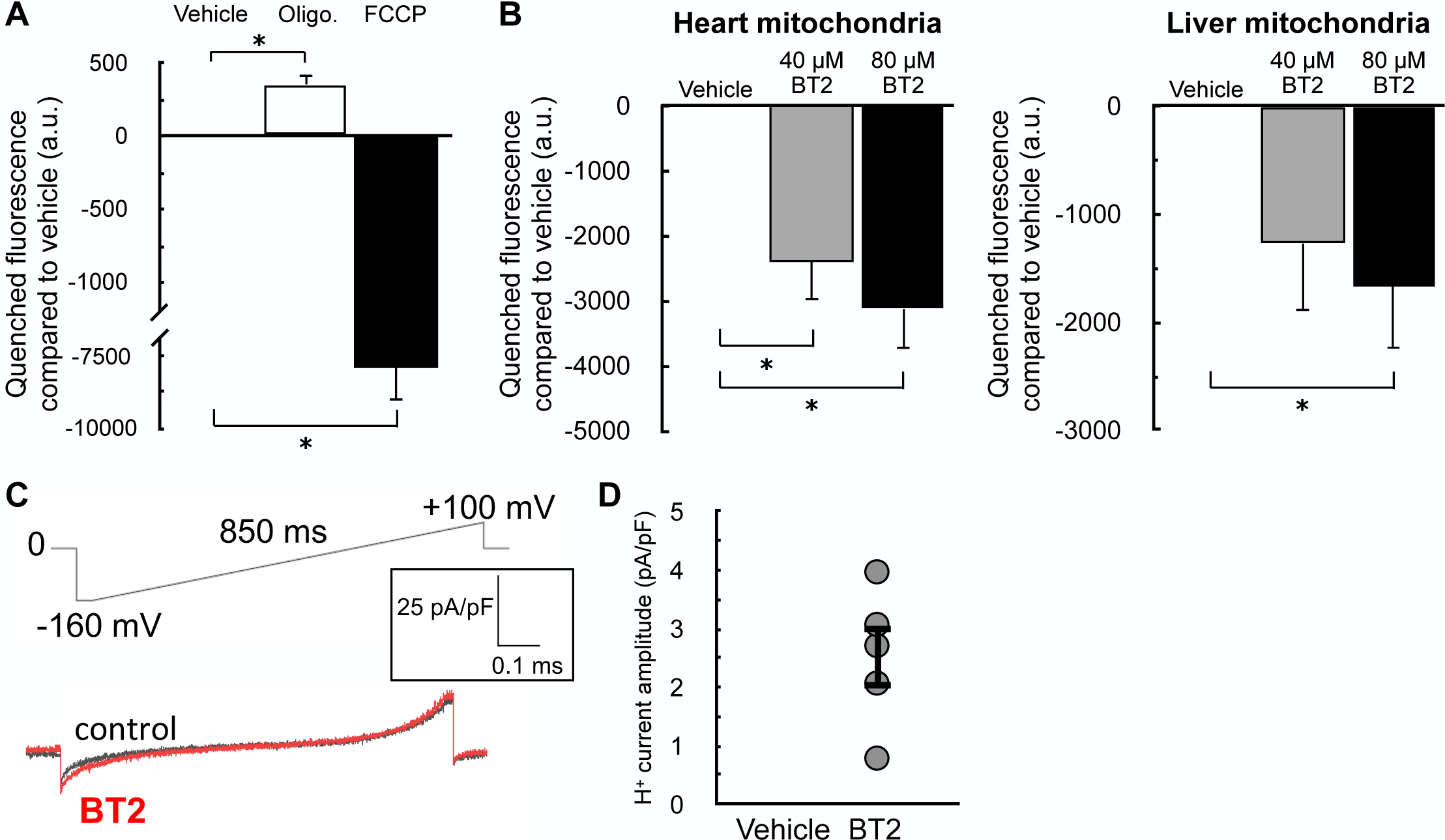
BT2 lowers membrane potential in isolated mitochondria and increases proton conductance across the mitochondrial inner membrane. (A) Quenched TMRE fluorescence in isolated rat liver mitochondria offered succinate, rotenone, and ADP as described in the Materials and Methods. Mitochondria were offered oligomycin (1 μg/mg mitochondrial protein) or FCCP (1μM) 5 min prior to measurements as controls to demonstrate the signal responds appropriately to known effector compounds. (n=3 biological replicates). (B) Quenched TMRE fluorescence as in (A) in response to 40μM BT2, 80μM BT2, or vehicle controls in isolated rat heart (*left*) or rat liver (*right*) mitochondria. (n=4 biological replicates). (C) Representative current induced by 100 µM BT2 in heart mitoplasts. (D) Current densities at −160 mV from (C). (n=5) All data are mean ± S.E.M. Statistical analysis was conducted with ANOVA followed by Dunnett’s post-hoc multiple comparisons tests. *, p < 0.05.

The acute increase in State 4_o_ respiration (Fig. 2) and decrease in membrane potential in isolated mitochondria strongly infer that BT2 increases proton conductance across the mitochondrial inner membrane [45]. However, a definitive, unassailable measure of mitochondrial uncoupling requires direct measurement of the proton current. We therefore conducted patch-clamp electrophysiological measurements of the inner membrane in mitoplasts derived from isolated heart mitochondria [48]. Indeed, a measurable increase in proton current could be observed across the inner membrane in response to BT2 (**Figs. 3C, D**). Taken together, results from respirometry, TMRE fluorescence, and membrane electrophysiology all point to a chemical uncoupling property of BT2 independent of its effects on BCAA metabolism.

### BT2 is ∼5-fold less potent than DNP

With clear evidence that BT2 can act as a chemical uncoupler, we then sought to gauge its potency in relation to the other well-known uncouplers DNP, FCCP, and Bam15. Similar to these chemical uncouplers, BT2 is lipophilic and contains a functional group that can reversibly dissociate with a proton **(Fig. 4A).** We therefore sought to quantify the potency of BT2 by measuring its concentration-response curve for State 4_o_ respiration alongside other chemical uncouplers in isolated liver mitochondria (**Figs. 4B, C**). Similar to DNP, BT2 is a far less potent uncoupler than either FCCP or Bam15, and requires >10^3^-fold more compound to elicit a similar change in proton leak-linked respiration (**Fig. 4D**). Relative to DNP, roughly 5 times the concentration of BT2 is required to elicit the same change in respiration in mitochondria isolated from both rat liver (**Fig. 4D**) and heart (**Fig. 4E**). Single-point measurements of quenched TMRE fluorescence also show a similar potency profile for BT2 relative to other uncouplers (**Fig. 4F**), as more BT2 is required to drop the mitochondrial membrane potential to a similar degree relative to other chemical uncouplers. Encouragingly, quantitative comparison of the proton current amplitude (pA/pF) elicited from 100 µM BT2 in electrophysiological studies (Fig. 3D) is 6-fold lower than that from 100 µM DNP [BT2: 2.52 ± 0.53 (S.E.M.; n=5); DNP: 15.82 ± 1.28 (S.E.M.; n=9) [32]]. Altogether, respiration and membrane potential measurements in isolated mitochondria show BT2 is three orders of magnitude less potent than Bam15 and FCCP, and broadly 5-fold milder than the well-studied and prototypical uncoupler DNP.

**Figure 4.**
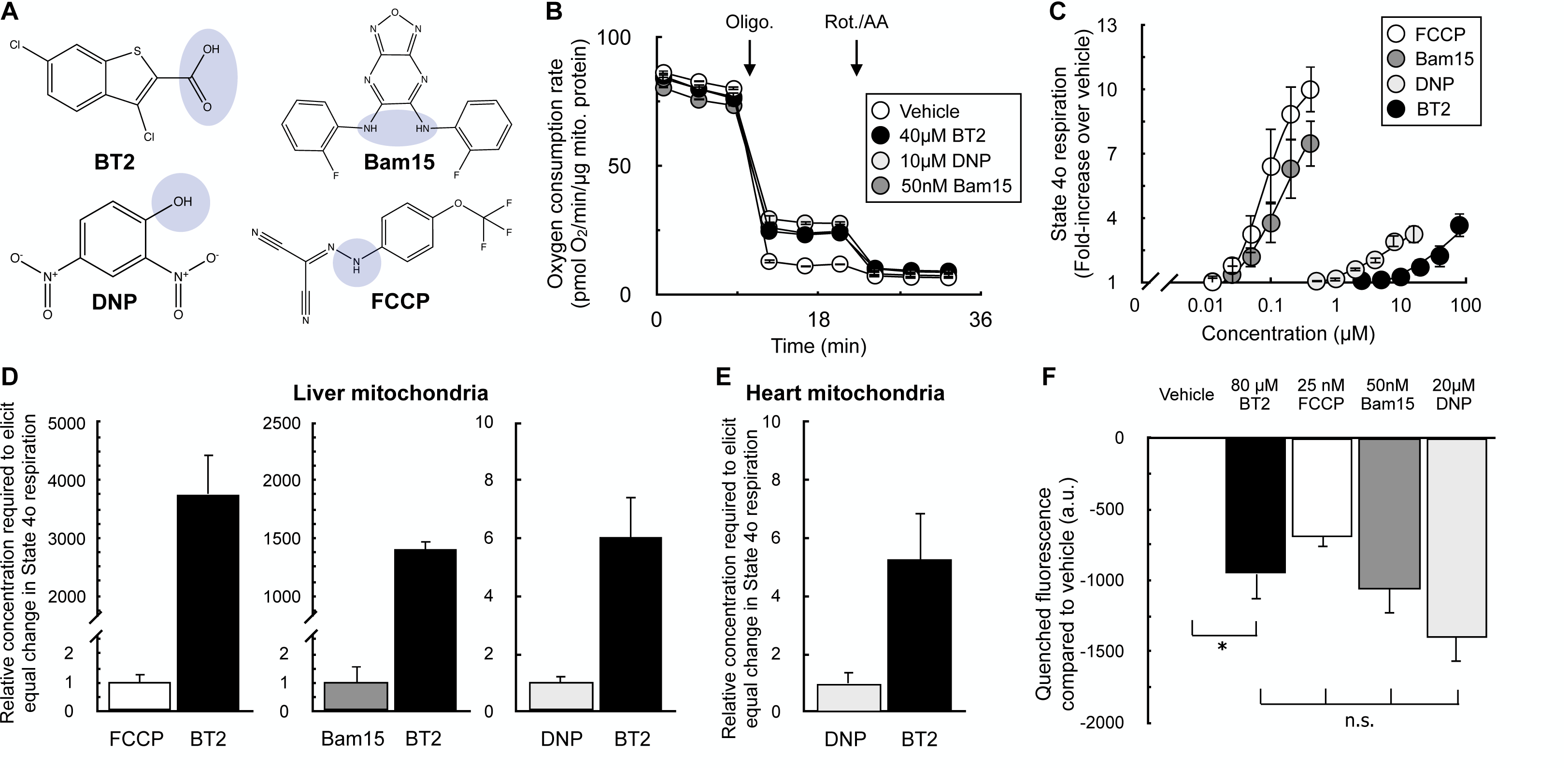
BT2 is a less potent chemical uncoupler than FCCP, Bam15, and DNP. (A) Chemical structures of BT2, DNP, Bam15, and FCCP with the functional group for weak acid/anion cycling highlighted in blue. (B) Representative trace of isolated liver mitochondria acutely offered 40μM BT2, 10μM DNP, or 50nM Bam15. Assay medium is supplemented with pyruvate, malate, and ADP as before. (n=4 technical replicates from a single experiment). (C) Aggregate fold-change in State 4_o_ respiration relative to vehicle controls from experiments conducted as in (B). [FCCP] and [Bam15] = 400 nM, 200 nM, 100 nM, 50 nM, 25 nM; [DNP] = 16 μM, 8 μM, 4 μM, 2 μM, 1 μM, 500 nM; [BT2] = 2.5, 5, 10, 20, 40, and 80μM. (n=4 biological replicates). (D) Data extrapolated from experiments conducted in (A) and (B) measuring difference in concentration required to elicit a two- and three-fold change in the State 4_o_ rate relative to vehicle controls. (n=4 biological replicates). (E) Values calculated as in (A) – (C) except with isolated rat heart mitochondria. (n=4 biological replicates). (F) Mitochondrial membrane potential measurements using quenched TMRE fluorescence with isolated liver mitochondria as in Figure 2 (A) – (C). (n=4 biological replicates). All data are mean ± S.E.M. Statistical analysis was conducted with ANOVA followed by Dunnett’s post-hoc multiple comparisons tests [(F)]. *, p < 0.05.

### BT2 uncouples via both AAC-dependent and AAC-independent mechanisms

Chemical uncouplers have at least two distinct mechanisms: (i) a conventional model whereby weak acid/anion cycling across the mitochondrial inner membrane dissipates the proton gradient across the inner membrane [49], and (ii) stimulating proton conductance through the ADP/ATP carrier (AAC) by binding to the nucleotide translocation site [32,50,51]. As such, we sought to determine whether BT2 also uncoupled via both AAC-dependent and AAC-independent mechanisms to provide further evidence that it is a conventional chemical uncoupler. We employed both pharmacologic and genetic strategies to inhibit AAC activity and measured BT2-stimulated increases in State 4_o_ respiration. In isolated rat liver mitochondria, inhibition of adenine nucleotide translocation with carboxyatractyloside (CATR) had a predictable effect of inhibiting ADP-stimulated respiration similarly to oligomycin (**Fig. 5A**). Quantifying the increase in State 4o respiration from BT2 in the presence and absence of CATR showed that AAC inhibition limited the chemical uncoupling independently of the substrate provided (**Figs. 5B, C**). To verify if the results could extend to intact cells, we quantified the effect of BT2 on respiration in C2C12 myoblasts with both *Slc25a4* (encoding AAC1) and *Slc25a5* (encoding AAC2) genetically ablated (‘double-knockout’ DKO) [32]. In both intact (**Fig. 5D-F**) and permeabilized cells (**Fig. 5F**), loss of *Aac1* and *Aac2* again resulted in the partial loss of BT2-stimulated uncoupling. In sum, results from both pharmacologic and genetic loss of AAC function point to BT2 having AAC-dependent and AAC-independent mechanisms of mitochondrial uncoupling similar to other well-characterized chemical uncouplers.

**Figure 5.**
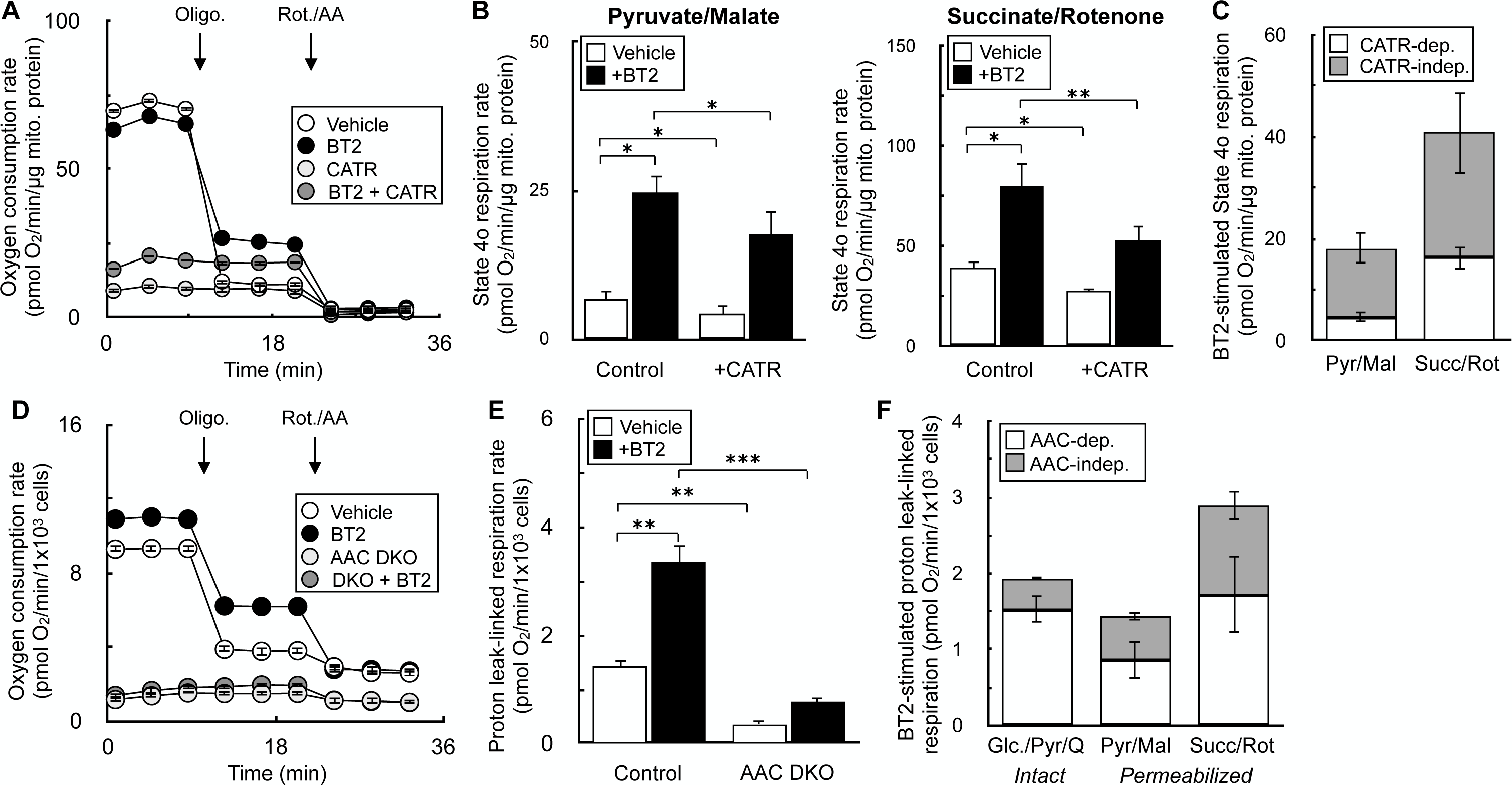
BT2 uncouples via both AAC-dependent and AAC-independent mechanisms. (A) Isolated rat liver mitochondria acutely offered 80μM BT2, 5μM carboxyatractyloside (CATR), both 80μM BT2 and 5μM CATR, or vehicle control 5 min prior to conducting measurements. Mitochondria were offered pyruvate, malate, and ADP as before. (n=10 technical replicates from a single experiment). (B) Collated State 4_o_ respiration from experiments as in (A) from rat liver mitochondria offered either pyruvate/malate (*left*) or succinate/rotenone (*right*). (n=4 biological replicates). (C) Collated State 4_o_ respiration from experiments as in (A) and (B) plotted as a stacked bar chart to gauge the proportion of CATR-sensitive (white) and CATR-insensitive (grey) uncoupling. (n=4 biological replicates). (D) Intact C2C12 myoblasts with or without genetic ablation of AAC1 and AAC2 (DKO) assayed in DMEM supplemented with glucose, pyruvate, and glutamine as described in the Materials and Methods. 80μM BT2 was offered acutely 5 min prior to conducting measurements. (n=10 technical replicates from a single experiment). (E) Collated proton leak-linked respiration from experiments using intact C2C12 myoblasts as in (D). (n=5 biological replicates). (F) Collated proton leak-linked respiration from experiments as in (D) and (E) plotted as a stacked bar chart to gauge the proportion of AAC-dependent (white) and AAC-independent (grey) uncoupling. Conditions are intact cells offered DMEM supplemented as in (A) (*left*) and permeabilized myoblasts offered either pyruvate/malate (*middle*) or succinate/rotenone (*right*). (n=5 biological replicates). All data are mean ± S.E.M. Statistical analysis was conducted with ANOVA followed by Dunnett’s post-hoc multiple comparisons tests. *, p < 0.05; **, p < 0.01; ***, p < 0.001.

### BT2 phenocopies DNP and lowers mitochondrial H_2_O_2_ efflux

Having demonstrated that BT2 can act as a chemical uncoupler distinct from its inhibition of the BCKDK, we next examined whether BT2 can reproduce hallmark biochemical and physiological responses resulting from chemical uncoupling. For example, it is well known that some forms of mitochondrial superoxide production are steeply dependent on the mitochondrial membrane potential [40,52]. As such, even mild chemical uncoupling that fractionally lowers the membrane potential can have a profound effect in lowering mitochondrial reactive oxygen species (ROS) production.

Previous work has shown that BT2 treatment reduces ROS production in cells ([21]). In *in vivo* pre-clinical heart failure models, BT2 administration preserves cardiac functions known to be sensitive to oxidative stress and protects from ROS-associated I/R and MI injury. ([12,20,21]). However, in these systems it is difficult to discriminate whether the beneficial effects of BT2 result from BCKDK inhibition or other uncharacterized mechanisms. We therefore tested whether BT2 could directly lower mitochondrial superoxide production derived from the respiratory chain.

Isolated rat heart mitochondria were offered succinate to generate superoxide resulting from a high proton motive force and reduced CoQ pool (i.e., “reverse electron transport”). Rates of superoxide production (measured as H_2_O_2_ efflux) were quantitatively comparable to previous reports [40], and the signal responds appropriately by reporting high rates of superoxide production in the presence of antimycin (**Fig. 6A, B**) [53]. As expected, BT2 significantly reduced mitochondrial H_2_O_2_ efflux. Importantly, the result could be phenocopied with the chemical uncouplers DNP and FCCP, each of which are structurally dissimilar and have no reported effects on the BCKDK. As such, the data indicate that mitochondrial uncoupling could be a mechanism by which BT2 can reduce mitochondrial ROS production independently of BCAA and BCKA metabolism.

**Figure 6.**
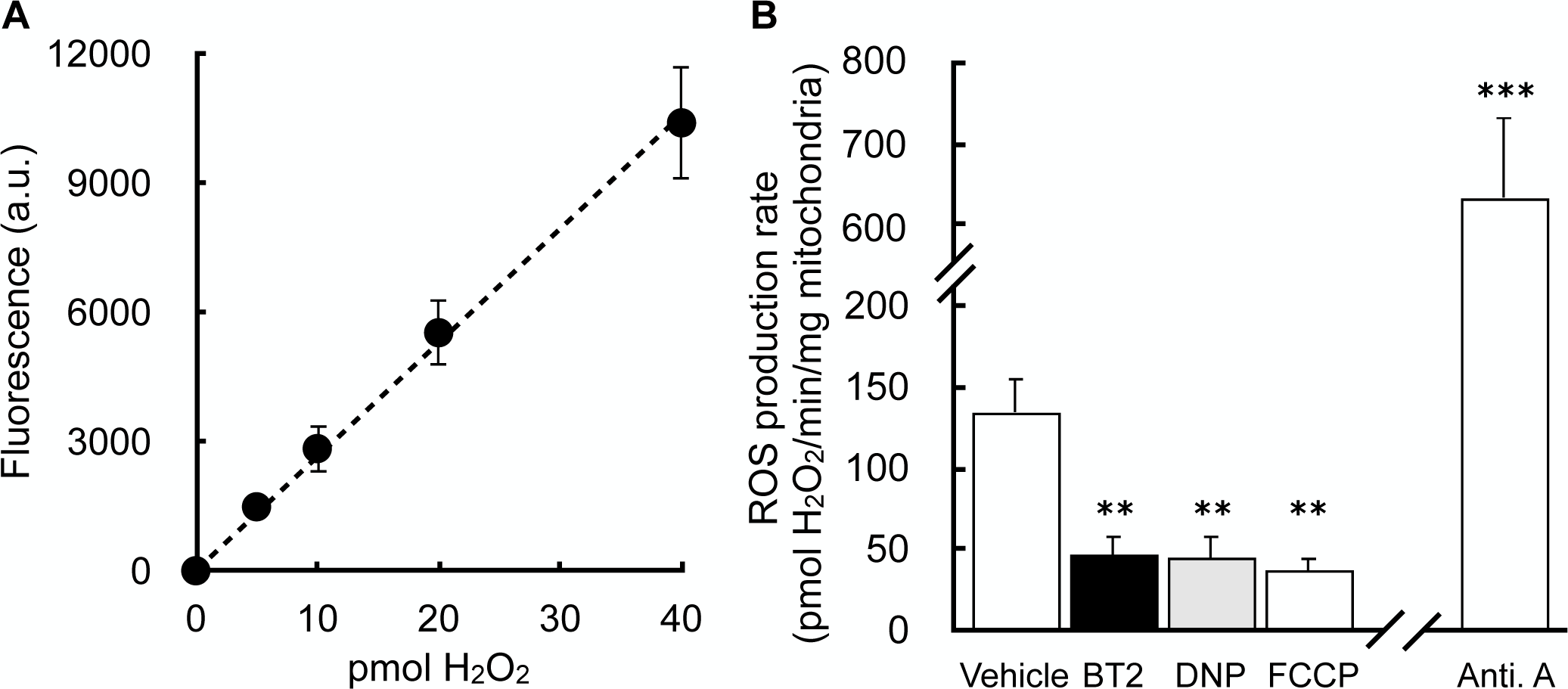
BT2 lowers mitochondrial H_2_O_2_ efflux. (A) Representative standard curve of exogenous H_2_O_2_ added on top of isolated mitochondria. (n=3 technical replicates from a single experiment). (B) H_2_O_2_ efflux in isolated mitochondria offered succinate in response to addition of 80μM BT2, 20μM DNP, 100nM FCCP, 1μM antimycin A, or vehicle control. (n=3-4 biological replicates). All data are mean ± S.E.M. Statistical analysis was conducted with ANOVA followed by Dunnett’s post-hoc multiple comparisons tests. **, p < 0.01; ***, p < 0.001.

### BT2 phenocopies DNP and reduces de novo lipogenesis

Although attenuation of mitochondrial ROS production by chemical uncoupling is a plausible mechanism for the cardioprotection afforded by BT2, it almost certainly cannot explain why BT2 protects from insulin resistance and hepatic steatosis. However, another hallmark response to chemical uncouplers is increased energy expenditure. Uncouplers create a ‘futile cycle’ where dissipation of the mitochondrial membrane potential stimulates respiratory chain activity [45,48]. This shifts the energy balance towards nutrient oxidation and away from lipid storage and accumulation. Decades of experimental evidence in rodents and humans support this mechanism [54–57], and chemical uncoupling is consistent with the myriad and sometimes rapid effects of BT2 in various metabolic disease models [26,58].

We examined whether BT2 could phenocopy DNP in increasing energy expenditure and reducing *de novo* lipogenesis in differentiated 3T3-L1 adipocytes. Indeed, acute treatment with either BT2 or DNP increased the basal oxygen consumption rate due to increased proton leak-linked respiration (**Fig. 7A, B**). We then conducted stable isotope tracing with uniformly labeled ^13^C_6_-glucose to further characterize any shared effects of BT2 and DNP. Both compounds also increased relative incorporation of glucose into the TCA cycle metabolites citrate and malate (**Fig. 7C**) and decreased the steady-state abundance of these and other TCA cycle intermediates (**Fig. 7D**), a profile consistent with an increased metabolic rate. Taken together, the respirometry and GC/MS data demonstrate that BT2 can increase energy expenditure via mitochondrial uncoupling.

**Figure 7.**
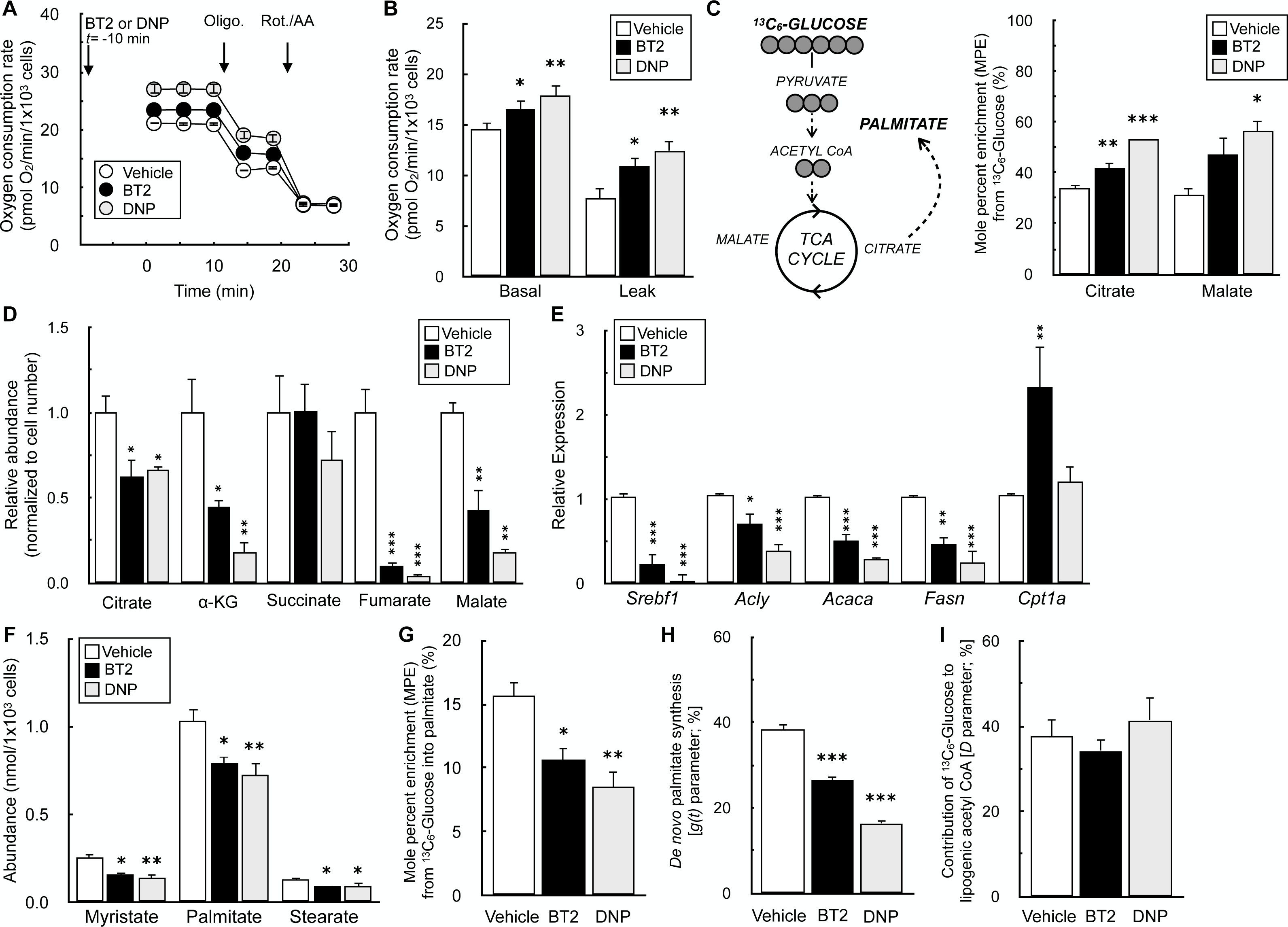
BT2 increases cellular energy expenditure and decreases *de novo* lipogenesis. (A) Representative oxygen consumption trace of differentiated 3T3-L1 adipocytes acutely offered 40μM BT2, or 10μM DNP 10 min. prior to initial measurements in DMEM supplemented with glucose, pyruvate, and glutamine as described in the Materials and Methods. (n=5 technical replicates from a single experiment). (B) Collated oxygen consumption rate parameters for 3T3-L1 cells as in (A). (n=5 biological replicates). (C) (*Left*) Simplified schematic of uniformly labeled ^13^C_6_-glucose enriching TCA cycle intermediates and palmitate. (*Right*) Mole percent enrichment (M.P.E.) from ^13^C_6_-glucose of TCA cycle intermediates citrate and malate in differentiated 3T3-L1 adipocytes treated for 72 hr. with 120μM BT2 or 40μM DNP in medium containing 2% serum. High concentrations of drug were used to account for drug binding to serum albumin. (n=3 biological replicates). (D) Relative metabolite abundances of TCA cycle intermediates, adjusted for cell number, for experiments conducted as in (C). (n=3 biological replicates). (E) qPCR analysis for genes associated with *de novo* lipogenesis as well as *Cpt1a* for cells treated as in (C). (n=5-8 biological replicates). (F) Abundances of fatty acids for experiments conducted as in (C). (n=3-4 biological replicates). (G) M.P.E. from ^13^C_6_-glucose into palmitate for experiments conducted as in (C). (n=5 biological replicates). (H) Rate of *de novo* lipogenesis [g(t) parameter] calculated by isotopomer spectral analysis (ISA) for experiments conducted as in (C). Averages provided by the model for each biological replicate are presented. Predicted g(t) values and 95% confidence intervals for each technical replicate are provided in Supplementary Table 1. (n=4 biological replicates). (I) Fractional contribution of glucose to the lipogenic acetyl CoA pool (‘D’ parameter) calculated by ISA. Averages provided by the model for each biological replicate are presented. Predicted ‘D’ values and 95% confidence intervals for each technical replicate are provided in Supplementary Table 1. (n=4 biological replicates). All data are mean ± S.E.M. Statistical analysis was conducted with ANOVA followed by Dunnett’s post-hoc multiple comparisons tests. *, p < 0.05; **, p < 0.01; ***, p < 0.001.

Lastly, we examined the effect of BT2 and DNP on *de novo* lipogenesis (DNL), reasoning that increased uncoupling would decrease lipid accumulation. Treatment of differentiated 3T3-L1 adipocytes for 72 hr. with either BT2 or DNP significantly reduced mRNA levels of both a master transcriptional regulator of lipid synthesis (*Srebpf1*) as well as individual enzymes involved in DNL (*Acly*, *Acaca*, *Fasn*) (**Fig. 7E**). Importantly, levels of *Cpt1a* – the gene encoding carnitine palmitoyltransferase-1a, which is rate-controlling for long chain fatty acid oxidation – were unchanged with DNP and increased with BT2. This result demonstrates that the reduction in DNL genes is not due to toxicity or other experimental artifacts, but rather due to a specific shift away from lipid synthesis.

Beyond gene expression, we validated that BT2 reduced DNL by GC/MS analysis of fatty acids. BT2 or DNP treatment for 72 hr. reduced steady state levels of myristate (14:0), palmitate (16:0), and stearate (18:0) (**Fig. 7F**). Furthermore, in contrast to increased labeling of TCA cycle intermediates (Fig. 7C), incorporation of ^13^C_6_-glucose into palmitate was decreased in response to administering either compound (**Fig. 7G**). To estimate the rate of palmitate synthesis, we applied isotopomer spectral analysis (ISA) to model the rate of *de novo* palmitate synthesis [g(t) parameter] and contribution of glucose-derived carbon to the lipogenic acetyl CoA pool (‘D’ parameter) [44]. As in other assays, BT2 and DNP showed qualitative similarities as both decreased the estimated rate of DNL but did not appreciably change the ‘D’ parameter (**Fig. 7H, I, Supplemental Table 1**). Overall, the gene expression, respiration, and mass spectrometry data show that BT2 phenocopies DNP in increasing energy expenditure and lowering *de novo* lipogenesis. Thus, the results establish chemical uncoupling as a putative mechanism to explain the therapeutic effects of BT2 on cardiometabolic disease (**Fig. 8**).

**Figure 8.**
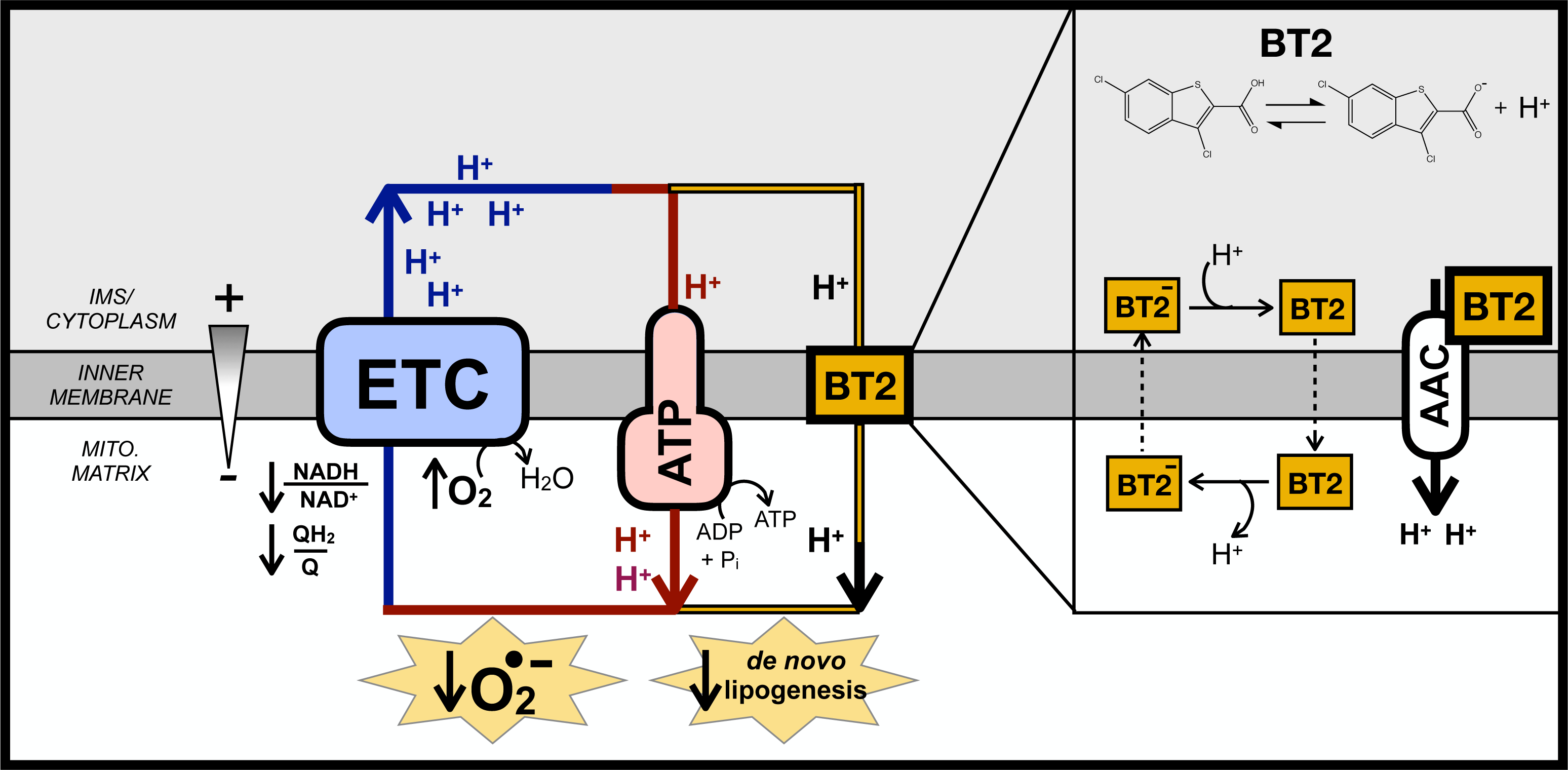
BT2 uncouples mitochondria to increase energy expenditure, lower superoxide production, and reduce *de novo* lipogenesis.

## DISCUSSION

Here we provide evidence that the BCKDK inhibitor BT2 is also a chemical uncoupler. Measurements of oxygen consumption, mitochondrial membrane potential, and H^+^ conductance all show BT2 consumes the membrane potential independently of ATP synthesis and uncouples mitochondria. BT2 stimulates proton conductance acutely (<5 minutes) and in reductionist systems bypassing BCKDK activity (i.e., isolated mitochondria offered various substrates as well as patch clamp electrophysiology of the mitochondrial inner membrane), demonstrating it is a bona fide chemical uncoupler independent of BCAA metabolism. Chemical uncoupling is a plausible, unifying mechanism for how BT2 can be therapeutic in various models of cardiovascular and metabolic disease independent of its effects on BCAA metabolism.

With respect to heart failure, it is generally accepted that excessive mitochondrial ROS production is an important pathological driver of cardiovascular disease [59–62]. A well-known property of uncouplers is limiting mitochondrial superoxide production by lowering the mitochondrial protonmotive force, thereby oxidizing the NADH/NAD^+^ and QH_2_/Q pools and lowering the probability of superoxide formation [52,63]. In addition to a direct effect on reducing mitochondrial ROS production from the respiratory chain, mitochondrial uncoupling is also likely to have an indirect effect in lowering ROS production by blocking induction of the permeability transition pore (PTP) [64–66]. The mitochondrial membrane potential provides the driving force for calcium uptake, and uncoupling mitigates the pathological matrix Ca^2+^ overload that triggers mitochondrial swelling and pathological opening of the PTP [66,67]. Importantly, both FCCP and DNP confer cardioprotection in *ex vivo* models of I/R injury using the Langendorff-perfused rat heart model [68,69], and heterologous expression of UCP1 in the mouse heart protects from ischemic damage and lowers markers of oxidative stress [70]. These proof-of-concept studies reinforce the notion that attenuation of ROS production via chemical uncoupling may explain why BT2 could be cardioprotective independent of its function as a BCKDK inhibitor. Moreover, it should be reinforced that given the steep dependence of ROS production and calcium uptake on the protonmotive force, even mild chemical uncoupling that fractionally lowers the membrane potential can have a substantive effect in lowering mitochondrial reactive oxygen species (ROS) production [40,52].

Regarding metabolic disease, it is well known that administration of chemical uncouplers and the resultant increase in energy expenditure can reduce insulin resistance, obesity, and hepatic steatosis in preclinical rodent models [54,71]. In fact, DNP was an effective anti-obesity treatment in the 1930s, but a poor safety profile and narrow therapeutic window – rather than a lack of efficacy – discontinued its clinical use [72,73]. Indeed, *in vivo* administration of BT2 in rodents qualitatively mimics the effect of other chemical uncouplers. In various preclinical models, BT2 is protective from glucose intolerance, insulin insensitivity, triglyceride accumulation, and hepatic inflammation similarly to DNP derivatives or Bam15 [22,24,26,29,58,74,75]. Several reports demonstrate BT2 administration does not change body weight [22,24,29]. However, this may be may be consistent with a mild uncoupler roughly five-fold less potent than DNP, and BT2 promotes a slight reduction in the respiratory exchange ratio in Zucker fatty rats [22].

The identification of a chemical uncoupling function for BT2 can readily explain some phenomena that are difficult to reconcile with a primary effect on BCAA metabolism. For example, previous reports show BT2 elicits a rapid euglycemic effect in glucose clamp studies less than an hour after administration [26]. Although this time frame is likely inconsistent with remodeling of BCAA metabolism, it is consistent with prior work showing acute effects on energy expenditure by DNP [58]. Moreover, our demonstration that DNP phenocopies BT2 in altering gene expression related to lipid metabolism suggests that chemical uncoupling could also explain why BT2 treatment blocks phosphorylation of acetyl CoA carboxylase at multiple sites in a manner not consistent with PP2Cm overexpression [22].

A critical aspect of our finding that BT2 is a chemical uncoupler is determining whether the *in vitro* concentrations required to stimulate uncoupling are relevant for *in vivo* studies. Our observation that BT2 concentrations as low as 2.5μM increase State 4o respiration raise the possibility that BT2 could elicit a mild uncoupling effect given that the concentrations achieved *in vivo* likely approach, if not surpass, the half-maximal inhibitory concentration for BCKDK inhibition (IC_50_ = 3-4μM) [23]. Peak circulating plasma concentrations of BT2 (∼1.0 mM for 40 mg/kg; ∼1.8 mM for 120 mg/kg) [76] and plasma protein binding measurements (99.3% protein bound) [23] suggest maximal, free BT2 concentrations reach single-digit micromolar concentrations and above. Moreover, pharmacokinetic (PK) data indicate a longer half-life in plasma relative to DNP and a clearance profile in plasma more similar to a controlled-release DNP analog [58,76]. Although tissue concentrations are a far more informative parameter, it is likely that BT2 would behave similarly to DNP as a lipophilic, membrane-targeting drug that can accumulate in tissues at concentrations at the same order of magnitude as circulating plasma levels [58,77]. Our demonstration that BT2 is roughly a five-fold weaker uncoupler than DNP may help explain why the drug is tolerable at doses that would be egregiously toxic for DNP. In total, available PK data support that chemical uncoupling should be a relevant, *in vivo* mechanism for BT2.

In addition to chemical uncoupling, it may be that other off-target effects of BT2 contribute to observed *in vitro* and *in vivo* phenotypes. For example, recent reports highlight shared pharmacology between angiotensin II type 1 receptor blockers (ARBs) and BT2 [78]. Encouragingly, regulation of the renin-angiotensin system may mechanistically explain the effect of BT2 on blood pressure despite *Bckdk* ablation [27]. Additional work suggests that BCKAs can inhibit the mitochondrial pyruvate carrier [79]. As BT2 acts as a branched-chain 2-oxoacid analog, it may also inhibit the MPC or other mitochondrial transporters or dehydrogenases. In fact, an inhibitory effect on pyruvate metabolism could explain our observation that BT2 stimulates *Cpt1a* expression in differentiated 3T3-L1 adipocytes while DNP does not [80].

In summary, our data suggest that chemical uncoupling by BT2 can explain many of the observed therapeutic benefits attributed to enhanced BCKDH activity, and encourage exploration of alternative hypotheses for existing data. For example, overexpression of the PP2Cm can reproduce some of the beneficial effects of BT2, but it is unclear to what extent this is due to BCKDH activity or other targets due to enzyme promiscuity [22,81]. It may also be that the insulin-sensitizing effects of sodium phenylbutyrate [25] – a lipophilic weak acid – are somewhat attributable to chemical uncoupling. Perhaps the strongest evidence against an alternative role for BT2 is the demonstration that the drug does not provide cardioprotection in whole-body *Bckdk*^-/-^ mice [27]. However, these animals are characterized by developmental impairments, neurological defects, and growth abnormalities that may warrant cautious interpretation of results [27,82]. Nonetheless, the data demonstrating that BT2 is a chemical uncoupler (i) reinforces the therapeutic promise of chemical uncoupling for a myriad of cardiovascular and metabolic diseases, and (ii) further highlights the importance of understanding to what extent BCAA accumulation is causative or associative in cardiovascular and metabolic disease pathogenesis.

## AUTHOR CONTRIBUTIONS

Conceptualization: AA, ASD, YW; Data curation: AA, AEJ, RT, CB, MW, AMB, ASD; Formal analysis: AA, AEJ, BD, RT, CB, MW, AMB, ASD; Funding acquisition: MW, YW, AMB, ASD; Investigation: AA, AEJ, BD, KPM, CB, LS, AMB, ASD; Methodology: AA, AEJ, BD, CB, LS, MW, AMB, ASD; Project administration: KR, OSS, YW, ASD; Resources: OSS, MW, YW, AMB, ASD; Supervision: AA, KR, OSS, YW, AMB, ASD; Writing - original draft: AA, ASD; and Writing - review & editing: AA, AEJ, BD, RT, KPM, CB, KR, OSS, LS, MW, YW, AMB, ASD.

## ABBREVIATIONS

AAC: ADP/ATP Carrier
BCAA: branched-chain amino acid
BCAT: branched-chain aminotransferase
BCKA: branched-chain α-ketoacid
BCKDH: branched-chain α-ketoacid dehydrogenase
BCKDK: branched-chain α-ketoacid dehydrogenase kinase
BT2: 3,6-dichlorobenzo[b]thiophene-2-carboxylic acid
CATR: carboxyatractyloside
DNP: 2,4-dinitrophenol
DNL: *de novo* lipogenesis
FCCP: Carbonyl cyanide 4-trifluoromethoxyphenylhydrazine
I/R: ischemia/reperfusion
MI: myocardial infarction
NRVM: neonatal rat ventricular myocytes
PP2Cm: mitochondrial protein phosphatase 2Cm
TMRE: tetramethylrhodamine, ethyl ester

## ACKNOWLEDGEMENTS

A.S.D. is supported by National Institutes of Health (NIH) Grants R35GM138003 and P30DK063491, as well as the W.M. Keck Foundation (995337) and the Agilent Early Career Professor Award. A.M.B. is supported by the National Institutes of Health (NIH) Grant R35GM143097 and the Pew Scholars in Biomedical Sciences. Y.W. is supported by Department of Defense CDMRP (PR191670). R.T and M.W are supported by an Ad Astra Fellowship. A.E.J. was supported by the UCLA Tumor Cell Biology Training Program (T32 CA009056).

---

**Supplemental Table 1 – ISA modeled values and 95% confidence intervals for individual technical replicates.**

