## Supplemental Table 1 for "The BCKDK inhibitor BT2 is a chemical uncoupler that lowers mitochondrial ROS production and *de novo* lipogenesis"

Supplemental Table 1 – ISA modeled values and 95% confidence intervals for individual technical replicates.

| Sample | Measurement | Value | Lower range | Upper range | Sample | Measurement | Value | Lower range | Upper range |
|---|---|---|---|---|---|---|---|---|---|
| **N1-NT1** | **D(M2-AcCoA)** | 0.309 | 0.264 | 0.353 | **N3-NT1** | **D(M2-AcCoA)** | 0.558 | 0.509 | 0.605 |
| | **D(M1-AcCoA)** | 0.027 | 0.006 | 0.052 | | **D(M1-AcCoA)** | 0.034 | 0.010 | 0.066 |
| | **1-D(AcCoA)** | 0.665 | 0.621 | 0.708 | | **1-D(AcCoA)** | 0.408 | 0.360 | 0.457 |
| | **g(t) palmitate** | 0.419 | 0.375 | 0.459 | | **g(t) palmitate** | 0.456 | 0.408 | 0.498 |
| | **D(TOTAL)** | 0.335 | 0.270 | 0.405 | | **D(TOTAL)** | 0.592 | 0.519 | 0.671 |
| **N1-NT2** | **D(M2-AcCoA)** | 0.288 | 0.240 | 0.335 | **N3-NT2** | **D(M2-AcCoA)** | 0.292 | 0.236 | 0.348 |
| | **D(M1-AcCoA)** | 0.027 | 0.006 | 0.055 | | **D(M1-AcCoA)** | 0.030 | 0.005 | 0.065 |
| | **1-D(AcCoA)** | 0.685 | 0.638 | 0.731 | | **1-D(AcCoA)** | 0.677 | 0.624 | 0.730 |
| | **g(t) palmitate** | 0.394 | 0.349 | 0.435 | | **g(t) palmitate** | 0.354 | 0.307 | 0.396 |
| | **D(TOTAL)** | 0.315 | 0.246 | 0.390 | | **D(TOTAL)** | 0.323 | 0.241 | 0.412 |
| **N1-NT3** | **D(M2-AcCoA)** | 0.244 | 0.194 | 0.291 | **N3-NT3** | **D(M2-AcCoA)** | 0.326 | 0.248 | 0.398 |
| | **D(M1-AcCoA)** | 0.032 | 0.009 | 0.064 | | **D(M1-AcCoA)** | 0.030 | 0.000 | 0.078 |
| | **1-D(AcCoA)** | 0.724 | 0.678 | 0.769 | | **1-D(AcCoA)** | 0.644 | 0.574 | 0.715 |
| | **g(t) palmitate** | 0.395 | 0.348 | 0.437 | | **g(t) palmitate** | 0.262 | 0.211 | 0.306 |
| | **D(TOTAL)** | 0.276 | 0.204 | 0.355 | | **D(TOTAL)** | 0.356 | 0.248 | 0.476 |
| **N1-BT2-1** | **D(M2-AcCoA)** | 0.322 | 0.255 | 0.385 | **N3-BT2-1** | **D(M2-AcCoA)** | 0.327 | 0.282 | 0.372 |
| | **D(M1-AcCoA)** | 0.029 | 0.000 | 0.070 | | **D(M1-AcCoA)** | 0.023 | 0.002 | 0.048 |
| | **1-D(AcCoA)** | 0.650 | 0.588 | 0.712 | | **1-D(AcCoA)** | 0.650 | 0.605 | 0.694 |
| | **g(t) palmitate** | 0.285 | 0.236 | 0.329 | | **g(t) palmitate** | 0.366 | 0.325 | 0.403 |
| | **D(TOTAL)** | 0.350 | 0.255 | 0.455 | | **D(TOTAL)** | 0.350 | 0.284 | 0.420 |
| **N1-BT2-2** | **D(M2-AcCoA)** | 0.299 | 0.221 | 0.370 | **N3-BT2-2** | **D(M2-AcCoA)** | 0.295 | 0.223 | 0.362 |
| | **D(M1-AcCoA)** | 0.030 | 0.000 | 0.079 | | **D(M1-AcCoA)** | 0.030 | 0.000 | 0.075 |
| | **1-D(AcCoA)** | 0.671 | 0.603 | 0.739 | | **1-D(AcCoA)** | 0.675 | 0.611 | 0.739 |
| | **g(t) palmitate** | 0.258 | 0.209 | 0.303 | | **g(t) palmitate** | 0.235 | 0.191 | 0.274 |
| | **D(TOTAL)** | 0.329 | 0.221 | 0.449 | | **D(TOTAL)** | 0.325 | 0.223 | 0.437 |
| **N1-BT2-3** | **D(M2-AcCoA)** | 0.231 | 0.076 | 0.320 | **N3-BT2-3** | **D(M2-AcCoA)** | 0.316 | 0.245 | 0.383 |
| | **D(M1-AcCoA)** | 0.046 | 0.004 | 0.169 | | **D(M1-AcCoA)** | 0.028 | 0.000 | 0.072 |
| | **1-D(AcCoA)** | 0.723 | 0.646 | 0.826 | | **1-D(AcCoA)** | 0.655 | 0.591 | 0.720 |
| | **g(t) palmitate** | 0.275 | 0.218 | 0.325 | | **g(t) palmitate** | 0.248 | 0.202 | 0.289 |
| | **D(TOTAL)** | 0.277 | 0.079 | 0.489 | | **D(TOTAL)** | 0.345 | 0.245 | 0.455 |
| **N1-DNP-1** | **D(M2-AcCoA)** | 0.355 | 0.236 | 0.455 | **N3-DNP-1** | **D(M2-AcCoA)** | 0.551 | 0.483 | 0.615 |
| | **D(M1-AcCoA)** | 0.024 | 0.000 | 0.095 | | **D(M1-AcCoA)** | 0.023 | 0.000 | 0.062 |
| | **1-D(AcCoA)** | 0.621 | 0.524 | 0.721 | | **1-D(AcCoA)** | 0.426 | 0.360 | 0.495 |
| | **g(t) palmitate** | 0.197 | 0.141 | 0.247 | | **g(t) palmitate** | 0.111 | 0.088 | 0.134 |
| | **D(TOTAL)** | 0.379 | 0.236 | 0.550 | | **D(TOTAL)** | 0.574 | 0.483 | 0.677 |
| **N1-DNP-2** | **D(M2-AcCoA)** | 0.387 | 0.243 | 0.505 | **N3-DNP-2** | **D(M2-AcCoA)** | 0.338 | 0.000 | 0.483 |
| | **D(M1-AcCoA)** | 0.022 | 0.000 | 0.105 | | **D(M1-AcCoA)** | 0.031 | 0.000 | 0.287 |
| | **1-D(AcCoA)** | 0.591 | 0.476 | 0.713 | | **1-D(AcCoA)** | 0.631 | 0.492 | 0.993 |
| | **g(t) palmitate** | 0.171 | 0.114 | 0.221 | | **g(t) palmitate** | 0.175 | 0.110 | 0.231 |
| | **D(TOTAL)** | 0.409 | 0.243 | 0.610 | | **D(TOTAL)** | 0.369 | 0.000 | 0.770 |
| **N1-DNP-3** | **D(M2-AcCoA)** | 0.346 | 0.201 | 0.456 | **N3-DNP-3** | **D(M2-AcCoA)** | 0.295 | 0.178 | 0.386 |
| | **D(M1-AcCoA)** | 0.025 | 0.000 | 0.109 | | **D(M1-AcCoA)** | 0.029 | 0.000 | 0.100 |
| | **1-D(AcCoA)** | 0.629 | 0.524 | 0.743 | | **1-D(AcCoA)** | 0.677 | 0.592 | 0.766 |
| | **g(t) palmitate** | 0.178 | 0.123 | 0.228 | | **g(t) palmitate** | 0.172 | 0.127 | 0.212 |
| | **D(TOTAL)** | 0.371 | 0.201 | 0.565 | | **D(TOTAL)** | 0.323 | 0.178 | 0.486 |
| **N2-NT1** | **D(M2-AcCoA)** | 0.224 | 0.187 | 0.260 | **N4-NT1** | **D(M2-AcCoA)** | 0.346 | 0.306 | 0.386 |
| | **D(M1-AcCoA)** | 0.031 | 0.012 | 0.054 | | **D(M1-AcCoA)** | 0.023 | 0.004 | 0.045 |
| | **1-D(AcCoA)** | 0.746 | 0.711 | 0.780 | | **1-D(AcCoA)** | 0.631 | 0.591 | 0.672 |
| | **g(t) palmitate** | 0.395 | 0.358 | 0.430 | | **g(t) palmitate** | 0.378 | 0.340 | 0.412 |
| | **D(TOTAL)** | 0.254 | 0.198 | 0.314 | | **D(TOTAL)** | 0.369 | 0.309 | 0.431 |
| **N2-NT2** | **D(M2-AcCoA)** | 0.387 | 0.337 | 0.436 | **N4-NT2** | **D(M2-AcCoA)** | 0.523 | 0.475 | 0.571 |
| | **D(M1-AcCoA)** | 0.024 | 0.002 | 0.052 | | **D(M1-AcCoA)** | 0.029 | 0.007 | 0.059 |
| | **1-D(AcCoA)** | 0.589 | 0.540 | 0.638 | | **1-D(AcCoA)** | 0.447 | 0.399 | 0.496 |
| | **g(t) palmitate** | 0.420 | 0.373 | 0.462 | | **g(t) palmitate** | 0.429 | 0.383 | 0.470 |
| | **D(TOTAL)** | 0.411 | 0.339 | 0.488 | | **D(TOTAL)** | 0.553 | 0.481 | 0.630 |
| **N2-NT3** | **D(M2-AcCoA)** | 0.249 | 0.203 | 0.293 | **N4-NT3** | **D(M2-AcCoA)** | 0.435 | 0.381 | 0.489 |
| | **D(M1-AcCoA)** | 0.030 | 0.008 | 0.059 | | **D(M1-AcCoA)** | 0.023 | 0.000 | 0.052 |
| | **1-D(AcCoA)** | 0.721 | 0.679 | 0.763 | | **1-D(AcCoA)** | 0.541 | 0.487 | 0.595 |
| | **g(t) palmitate** | 0.393 | 0.350 | 0.432 | | **g(t) palmitate** | 0.412 | 0.363 | 0.455 |
| | **D(TOTAL)** | 0.279 | 0.212 | 0.352 | | **D(TOTAL)** | 0.459 | 0.381 | 0.541 |
| **N2-BT2-1** | **D(M2-AcCoA)** | 0.398 | 0.295 | 0.492 | **N4-BT2-1** | **D(M2-AcCoA)** | 0.300 | 0.246 | 0.352 |
| | **D(M1-AcCoA)** | 0.029 | 0.000 | 0.094 | | **D(M1-AcCoA)** | 0.026 | 0.001 | 0.058 |
| | **1-D(AcCoA)** | 0.573 | 0.480 | 0.669 | | **1-D(AcCoA)** | 0.674 | 0.624 | 0.724 |
| | **g(t) palmitate** | 0.231 | 0.172 | 0.281 | | **g(t) palmitate** | 0.266 | 0.226 | 0.302 |
| | **D(TOTAL)** | 0.427 | 0.295 | 0.586 | | **D(TOTAL)** | 0.326 | 0.247 | 0.411 |
| **N2-BT2-2** | **D(M2-AcCoA)** | 0.282 | 0.203 | 0.354 | **N4-BT2-2** | **D(M2-AcCoA)** | 0.317 | 0.266 | 0.367 |
| | **D(M1-AcCoA)** | 0.032 | 0.000 | 0.084 | | **D(M1-AcCoA)** | 0.023 | 0.000 | 0.053 |
| | **1-D(AcCoA)** | 0.685 | 0.619 | 0.753 | | **1-D(AcCoA)** | 0.660 | 0.611 | 0.709 |
| | **g(t) palmitate** | 0.279 | 0.227 | 0.325 | | **g(t) palmitate** | 0.232 | 0.198 | 0.263 |
| | **D(TOTAL)** | 0.315 | 0.203 | 0.437 | | **D(TOTAL)** | 0.340 | 0.266 | 0.420 |
| **N2-BT2-3** | **D(M2-AcCoA)** | 0.369 | 0.271 | 0.459 | **N4-DNP-1** | **D(M2-AcCoA)** | 0.365 | 0.247 | 0.468 |
| | **D(M1-AcCoA)** | 0.030 | 0.000 | 0.091 | | **D(M1-AcCoA)** | 0.026 | 0.000 | 0.093 |
| | **1-D(AcCoA)** | 0.601 | 0.514 | 0.690 | | **1-D(AcCoA)** | 0.609 | 0.508 | 0.714 |
| | **g(t) palmitate** | 0.252 | 0.193 | 0.303 | | **g(t) palmitate** | 0.183 | 0.133 | 0.229 |
| | **D(TOTAL)** | 0.399 | 0.271 | 0.549 | | **D(TOTAL)** | 0.391 | 0.247 | 0.561 |
| **N2-DNP-1** | **D(M2-AcCoA)** | 0.464 | 0.319 | 0.590 | **N4-DNP-2** | **D(M2-AcCoA)** | 0.340 | 0.271 | 0.408 |
| | **D(M1-AcCoA)** | 0.021 | 0.000 | 0.117 | | **D(M1-AcCoA)** | 0.019 | 0.000 | 0.056 |
| | **1-D(AcCoA)** | 0.515 | 0.390 | 0.650 | | **1-D(AcCoA)** | 0.641 | 0.576 | 0.706 |
| | **g(t) palmitate** | 0.150 | 0.094 | 0.201 | | **g(t) palmitate** | 0.163 | 0.129 | 0.195 |
| | **D(TOTAL)** | 0.485 | 0.319 | 0.707 | | **D(TOTAL)** | 0.359 | 0.271 | 0.464 |
| | | | | | **N4-DNP-3** | **D(M2-AcCoA)** | 0.316 | 0.239 | 0.388 |
| | | | | | | **D(M1-AcCoA)** | 0.021 | 0.000 | 0.064 |
| | | | | | | **1-D(AcCoA)** | 0.663 | 0.594 | 0.732 |
| | | | | | | **g(t) palmitate** | 0.162 | 0.126 | 0.194 |
| | | | | | | **D(TOTAL)** | 0.337 | 0.239 | 0.452 |
